# An efficient extension of N-mixture models for multi-species abundance estimation

**DOI:** 10.1101/073577

**Authors:** Juan Pablo Gomez, Scott K. Robinson, Jason K. Blackburn, José Miguel Ponciano

## Abstract

1. In this study we propose an extension of the N-mixture family of models that targets an improvement of the statistical properties of rare species abundance estimators when sample sizes are low, yet typical size for tropical studies. The proposed method harnesses information from other species in an ecological community to correct each species’ estimator. We provide guidance to determine the sample size required to estimate accurately the abundance of rare tropical species when attempting to estimate the abundance of single species.

2. We evaluate the proposed methods using an assumption of 50-m radius plots and perform simulations comprising a broad range of sample sizes, true abundances and detectability values and a complex data generating process. The extension of the N-mixture model is achieved by assuming that the detection probabilities of a set of species are all drawn at random from a beta distribution in a multi-species fashion. This hierarchical model avoids having to specify a single detection probability parameter per species in the targeted community. Parameter estimation is done via Maximum Likelihood.

3. We compared our multi-species approach with previously proposed multi-species N-mixture models, which we show are biased when the true densities of species in the community are less than seven individuals per 100-ha. The beta N-mixture model proposed here outperforms the traditional Multi-species N-mixture model by allowing the estimation of organisms at lower densities and controlling the bias in the estimation.

4. We illustrate how our methodology can be used to suggest sample sizes required to estimate the abundance of organisms, when these are either rare, common or abundant. When the interest is full communities, we show how the multi-species approaches, and in particular our beta model and estimation methodology, can be used as a practical solution to estimate organism densities from rapid inventory datasets. The statistical inferences done with our model via Maximum Likelihood can also be used to group species in a community according to their detectabilities.

## 1 Introduction

Unbiased abundance and occupancy estimates are of paramount value for making inferences about ecological processes and making sound conservation decisions (Hubbell, 2001; Leibold *et al*., 2004; Margules & Pressey, 2000). To date, quantitative ecologists have proposed several statistical methods to estimate species’ detection probabilities and use these to correct occupancy or abundance estimates (Denes *et al*., 2015). Our study was motivated by the attempt to use these novel models to estimate the abundance of rare species in tropical communities. In these communities, it is well-known that abundance distributions are typically characterized by long right tails with few abundant species and many rare ones (Hubbell, 2001; Stratford & Robinson, 2005). Such high proportion of rare species in the overall community makes it very difficult to obtain enough detections during field surveys for appropriate estimation of both abundance and detection probability for many, if not the majority of species. When we extensively tested via simulations these recent methodologies, we found persistent bias in estimates of low abundances that corresponded to abundance ranges previously not dealt with in temperate forest studies yet common in neotropical studies (see also Yamaura, 2013; Yamaura *et al*., 2016). As an answer to this problem, here we present an alternative, community-based abundance estimation approach that markedly improves these estimates. Our method is widely applicable in communities marked by patterns of rare abundance (Stratford & Robinson, 2005; Robinson *et al*., 2000) or other ecological systems characterized by rare events (e.g. Seabloom *et al*., 2015).

In the single-species N-mixture, the model is used to estimate the abundance given imperfect detection (MacKenzie *et al*., 2002; Martin *et al*., 2005; Royle & Dorazio, 2008). It uses spatially and temporally replicated counts in which the counts of species *y* are binomially distributed with *N* being the total number of individuals available for detection and *p* the probability of detecting an individual of that species (Royle, 2004). The model is hierarchical because the abundance *N* is assumed to be a latent (i.e., unobserved), random process adopting a discrete probability distribution (*e.g*., Poisson). Inferences about the abundance of the species of interest therefore rely on estimating the detection probability and the underlying parameters of the distribution giving rise to *N* (Royle, 2004). Alternatively, multi-species models have been proposed to deal with estimating the abundance and occupancy of species with a limited amount of detections (see Iknayan *et al*., 2014; Denes *et al*., 2015, for reviews). These models have the advantage of “borrowing information” from abundant species in the community to estimate parameters of rare ones (Zipkin *et al*., 2009; Ovaskainen & Soininen, 2011; Yamaura *et al*., 2016, 2011; Chandler *et al*., 2013; Barnagaud *et al*., 2014). Most of the research and advances in the proposition of multi-species models has focused on estimating occupancy (Iknayan *et al*., 2014; Denes *et al*., 2015), even though understanding the abundance and rarity of species is one of the main goals of ecology (Yamaura *et al*., 2016; Hubbell, 2001; McGill *et al*., 2007).

In recent multi-species abundance models, both abundance and detection probabilities are assumed to be normally distributed random effects at the logit or log scales governed by a community’s “hyper-parameters” (Iknayan *et al*., 2014). For these reasons, they have been named community abundance models because they focus on describing the characteristics of the entire community from spatially and temporally replicated counts or detections (Yamaura *et al*., 2011, 2012, 2016). The main assumption behind the community abundance models is that groups of species in the community might share characteristics that make their abundance and detection probability likely to be correlated (Yamaura *et al*., 2011, 2012, 2016; Sauer & Link, 2002; Barnagaud *et al*., 2014; Ruiz-Gutiérrez *et al*., 2010). These types of abundance community models have been useful for estimating diversity properties of species assemblages while accounting for imperfect detection (Yamaura *et al*., 2011, 2012).

While the assumption of normally distributed logit-transformed random effects for detection probabilities of species across the community is statistically convenient, other probability distributions might have properties more directly related. Martin *et al*. (2011), for example, proposed a single-species abundance estimation model that allowed individuals within a species to vary in detection probability. They assumed that detection probabilities in a species were described by a beta distribution that naturally ranges between [0-1]. The latter assumption is convenient for community abundance models as well, because it eliminates the need of the logit transformation. Furthermore, Dorazio *et al*. (2013) showed that the beta distribution can be parametrized to reflect the mean detection probability among species and their degree of similarity making the two parameters that determine the shape of the beta distribution ecologically interpretable.

In this study, we: (1) increase the simulation scenarios presented in Yamaura (2013) to provide a full baseline for the sampling design for ecologists who want to estimate the abundance of tropical organisms (or any system with rare occurrence or detection difficulties) using N-mixture models, (2) propose an alternative multi-species abundance model that uses a beta distribution for the random effects of detection probability instead of a normal distribution, (3) propose a maximum likelihood approach for multi-species abundance estimation using Data Cloning and (4) compare our alternative multi-species abundance model to one previously proposed. Our study focuses on scenarios in which species have already been detected but the number of detections per species are insufficient to estimate detection-corrected abundances (i.e., low-abundance species). Our study does not focus on estimating the number or identity of unseen species. Instead we point to alternative models developed to account for this type of uncertainty (e.g. Dorazio & Royle, 2005; Royle & Dorazio, 2008; Tingley & Beissinger, 2013).

### 1.1 The Model

In the following section, after summarizing the widely used N-mixture models, we develop a multi-species model extension that allows a more accurate estimation of the abundance of rare species. Our approach differs from other multi-species abundance estimation by assuming that detection probabilities in a community are the product of a beta distribution instead of a logit transformation of normally distributed random effects.

According to an N-mixture model coded for one species, we let *y_ij_* be the number of individuals for that species in the *i*^th^ spatially replicated sampling unit and *j*^th^ temporal replicate of the sampling unit. Let *p* be the individual detection probability for that species. Finally, let *n_i_* be the fixed number of individuals available for detection in the i^th^ sampling unit. If we assume that the counts are binomially distributed, the likelihood of the counts (*y_ij_*) for a given species is

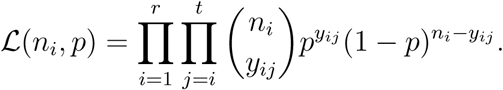
for *i* = 1, 2, 3 … *r* and *j* = 1, 2, 3 … *t*, where *r* is the total number of spatial replicates sampled and *t* is the number of times each spatial replicate was visited (Royle, 2004). In bird studies, for example, a common method used to survey individual populations or communities is fixed-radius plots (Hutto *et al*., 1986; Bibby *et al*., 2000). In this case, the researcher randomly locates 50-meter radius spatially replicated plots across the study area that are visited at different times. From here on, we will make our assumptions and definitions around this scenario in which 50-meter plots refer to spatial replicates of the sampling area and visits refers to temporal replicates of the count process in each plot. Also, in accord with conventions from bird literature, we will name each 50-m radius plot as a point count.

The N-mixture model assumes that the number of individuals available for detection in a point count is in fact unknown and random. Thus, this number is considered to be a latent variable, modeled with a Poisson process with mean λ. In what follows, λ is defined as the mean number of individuals per unit area, and we will refer to it as the “density”. We will write *N_i_* ∼ Pois (λ), where we have used the convention that lowercase letters such as *n_i_* denote a particular realization of the (capitalized) random variable *N_i_*. We note in passing that matrices will also be denoted with a capital letter, but will be written in bold. To compute the likelihood function, one then has to integrate the binomial likelihood over all the possible realizations of the Poisson process,

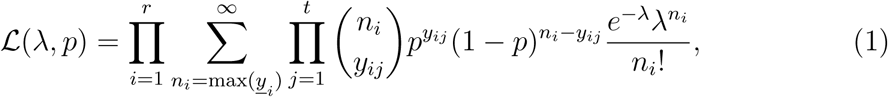
where *<u>y</u>_i_* is a vector of length *r* with the observed counts for that species for *i*^th^ point count. If the objective is to estimate the abundance of *S* species, the overall likelihood is simply written as the product of all the individual species’ likelihoods, *i.e*.,

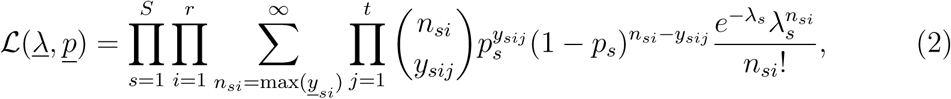
where *<u>y</u>_si_* is a vector of length *r* with the observed counts for species *s* in the *i*^th^ point count, and both <u>λ</u> = (λ_1_,…, λ*_S_*} and *<u>p</u>* = {*p*_1_,…, *p_S_*} are vectors of length *S*. To avoid the proliferation of parameters one could assume that all the *p_s_, s* = 1,…,*S* come from a single probability model that describes the community-wide distribution of detection probabilities (Yamaura *et al*., 2011, 2012, 2016; Sauer & Link, 2002; Barnagaud *et al*., 2014; Ruiz-Gutiérrez *et al*., 2010). In this case, each species’ detection probability can be modeled with a beta distribution. Let *P*_1_, *P*_2_, …, *P_S_* ∼ Beta(*α, β*). The probability density function of the random detection probabilities is then 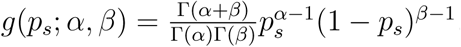.

Following Dorazio *et al*. (2013), we parameterize the Beta distribution as 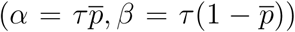 such that the parameters are related to biological processes. Here, 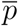 is the mean detection probability among species in the community and τ is a measurement of the similarity in detection probabilities (precision parameter; Dorazio *et al*., 2013). Note that 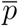 is equivalent to μ in Dorazio *et al*. (2013) parametrization but we avoid the use of μ in this proposition to avoid confusions with alternative models presented below. In this parametrization, the expected value and variance of P are given by 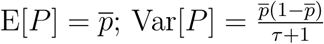.

The overall likelihood function now integrates over all the realizations of the community-wide detection probabilities *P_s_:*

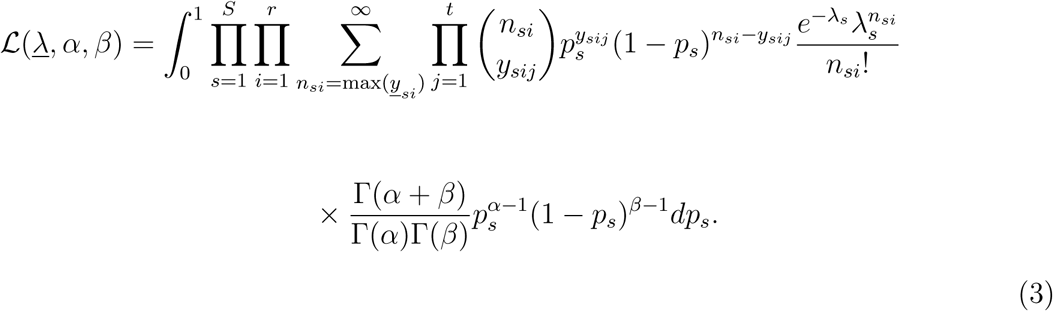

The usefulness of specifying the likelihood in this way is that in the case in which many species are rare, we can use the information on the abundant species to estimate the detection probability, leaving the actual counts to estimate only the abundance of the species. Note that by integrating the beta process at the outmost layer of the model, we are following the sampling structure. When this approach is used and the integral is tractable, the resulting distribution is a multivariate distribution with a specific covariance structure (Sibuya *et al*., 1964). Thus, we expect our approach to result in a multivariate distribution of counts with a covariance structure arising naturally from the sampling design and the assumed underlying beta process of detectabilities (see Table 1 for further description of the Beta N-mixture model).

**Table 1:**
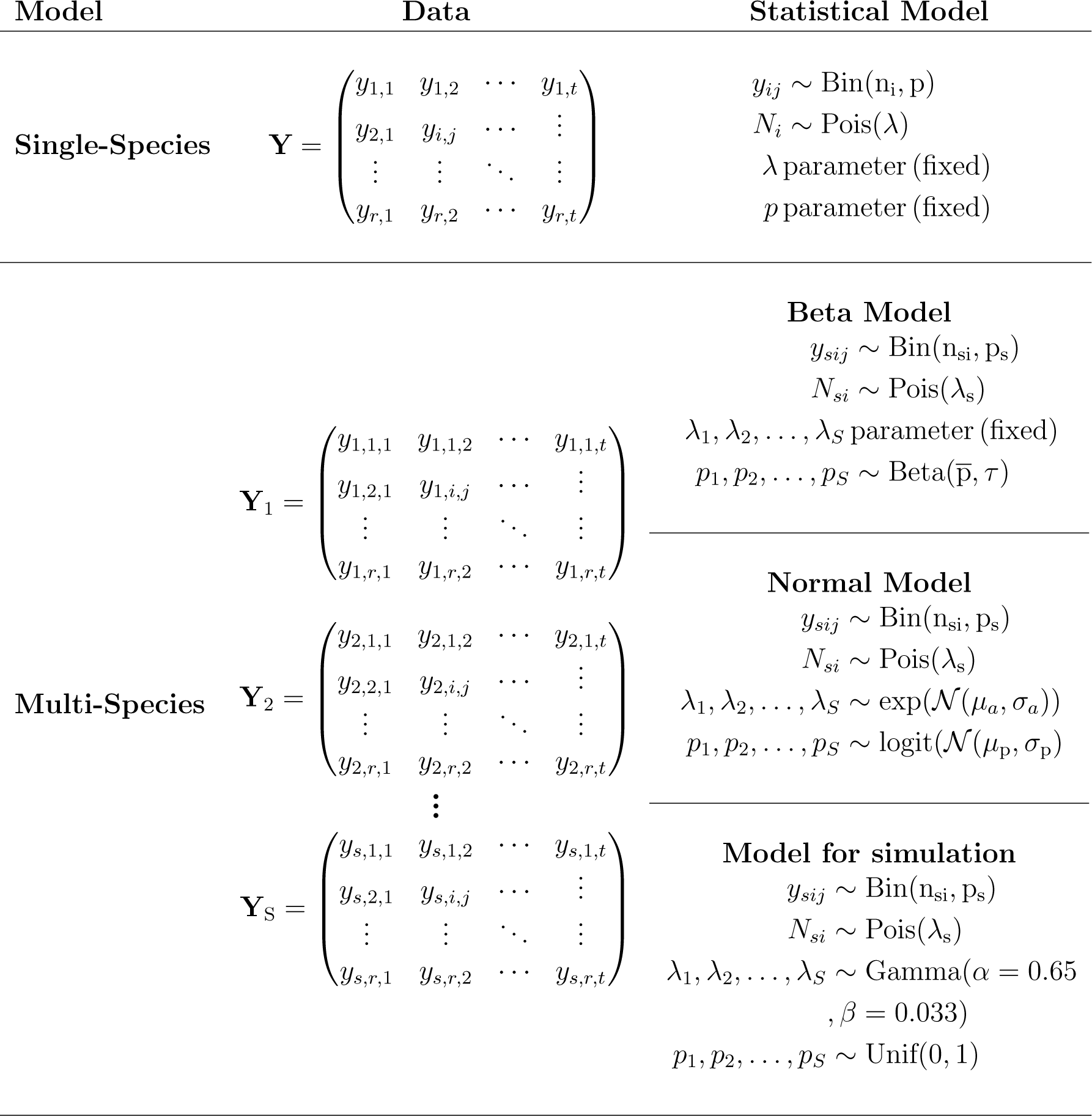
Summary of single and multi-species models used in this study. We also describe the model used to generate the simulated data for comparison between the multi-species models. *y* represents the observed counts, *N* the random variable of unobserved number of individuals *n* available for detection in plot *i, p* the detection probability, and λ the density of species *s*.

### 1.2 Maximum Likelihood Estimation

One drawback of the beta-N-mixture, and other models for multi-species abundance estimation, is their computational complexity, which imposes a substantial numerical challenge for Maximum Likelihood (ML) estimation. This problem is not unique to abundance estimation as it occurs in many other hierarchical models in ecology (Lele & Dennis, 2009). For these reasons, parameter estimation in hierarchical models is usually performed under a Bayesian framework (Cressie *et al*., 2009). To date, however, many numerical approximations for obtaining the Maximum Likelihood Estimates (MLEs) for hierarchical models have been proposed (de Valpine, 2012). The “Data Cloning” (DC) methodology has proven to be a reliable approach to obtaining MLEs, testing hypotheses, model selection, and unequivocally measuring the estimability of parameters for hierarchical models (Lele *et al*., 2010; Ponciano *et al*., 2012). The method proposed by Lele *et al*. (2007, 2010) uses the Bayesian computational approach coupled with Monte Carlo Markov Chain (MCMC) to compute MLEs of parameters of hierarchical models and their asymptotic variance estimates (Lele *et al*., 2007). The DC protocol is advantageous as one only needs to compute means and variances of certain posterior distributions.

Data Cloning proceeds by performing a typical Bayesian analysis on a dataset that consists of *k* copies of the originally observed data set. In other words, to implement this method, one has to write the likelihood function of the data as if one had observed k identical copies of the data set. Then, Lele *et al*. (2007, 2010) showed that as k grows large, the mean of the resulting posterior distribution converges on the MLE. In addition, for continuous parameters such as 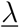, 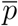 and τ, the variance covariance matrix of the posterior distribution converges to 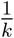 times the inverse of the observed Fisher’s information matrix. Thus, the variance estimated by the posterior distribution can be used to calculate Wald-type confidence intervals of the parameters (Lele *et al*., 2007, 2010). The advantage of DC over traditional Bayesian algorithms is that while in Bayesian algorithms the prior distribution might have influence over the posterior distribution, in DC the choice of the prior distribution does not determine the resulting estimates. In our case, the hierarchical statistical model for every species *s* in *s* = 1, 2, …, *S* is

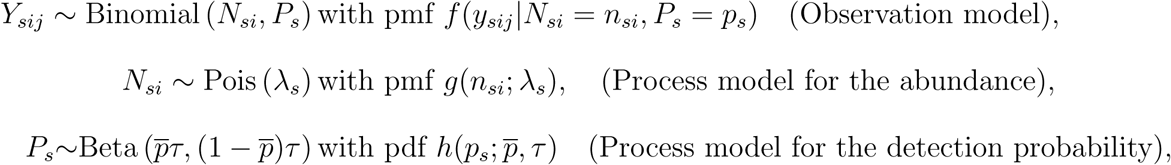
where *s* = 1, 2,…, *S, i* = 1,2,…, *r* and *j* = 1, 2,…, *t* and pmf and pdf correspond to the probability mass function and probability density functions respectively. According to our model, the values of λ_1_, λ_2_,…, λ*_S_* are parameters to be estimated. MLE of our model parameters would then generate point estimates of these parameters. In a Bayesian framework, however, parameters are random variables. Accordingly, the values of 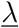, 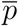 and τ would be modeled as random variables themselves that have a posterior distribution 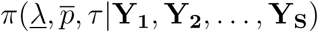. The Bayesian point estimates would typically be taken to be the posterior means or modes (although in a pure Bayesian approach the object of inference is the entire posterior distribution). We mention this Bayesian approach because, as we describe above, the DC methodology “tricks” a Bayesian estimation setting into yielding the MLEs. For this model, the specification of the Bayesian approach would require sampling from the following posterior distribution:

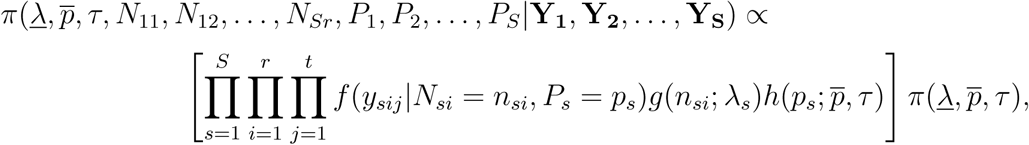
where 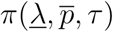 is the joint prior of the model parameters. Samples from an MCMC of this posterior distribution would yield many samples of the parameters

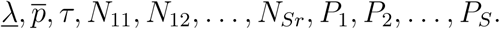
In order to sample from the marginal posterior 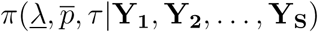 one only needs to look at the samples of the subset of 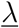, 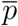 and, τ. The DC approach proceeds similarly, except one needs to sample from the following posterior distribution:

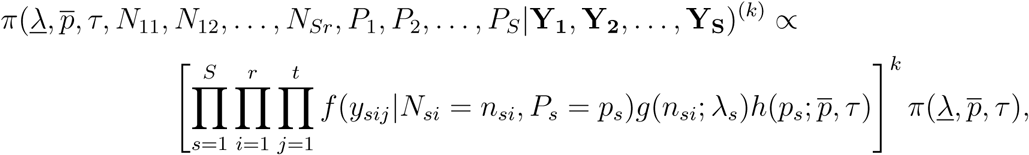

The notation ^(k)^ on the left side of this equation does not denote an exponent but the number of times the data set was “cloned”. On the right hand side, however, *k* is an exponent of the likelihood function based on the original data (*i.e*. un-cloned data; 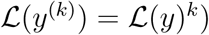. The MLEs of 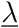, 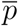 and, τ are then simply obtained as the empirical average of the posterior distribution 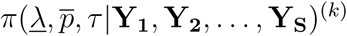 and the variance of the estimates are given by 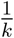 times the variance of this posterior distribution.

## 2 Methods

### 2.1 Estimation for Single Species

To determine the minimum sample size required for accurate estimation of the abundance of tropical species, we used a series of simulations in which we varied the number of point counts (*r*), visits to point counts (*t*; 50 meter fixed radius), density (mean number of individuals) in a 100 ha plot (λ), and detection probability (*p*). Point counts were assumed to be randomly located in a 100-ha plot. We varied *r* between 5 and 50, *t* between 2 and 20, λ = 1, 2, 3, 4, 5, 7, 10, 15, 25, 40, 55, 65, 75, 85, 100 and *p* between 0.1 and 0.9. Even though we assumed that λ was at a scale of individuals/100ha, because of the sampling area and design, the actual estimates are in individuals/0.78-ha. Thus, in this section and throughout out the rest of the sections, we estimated λ = *individuals*/0.78 − ha and extrapolate the estimates by applying 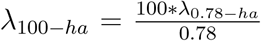. For the latter, λ_100-ha_ represents the density of an individual species *s* in a 100-ha plot and λ_0_._78-ha_ represents the density of a species in a point count with area of 0.78 ha. The area of the point counts corresponds with the area of a 50-m radius circular plot calculated as π * 50^2^ = 7854m^2^ ≈ 0.78ha. For each combination of parameters, we simulated 170 data sets and estimated λ_0_._78-_*_ha_* and *p* using equation 1. In each simulation, we computed the relative bias of the abundance estimate by using, bias 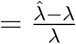, where 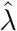 is the MLE for a particular data set and λ is the true value of the parameter. Finally, we retained the mean bias for each combination of model parameters. We considered an acceptable bias to be lower than 0.1, which is a 10% difference between the estimate and the true population density. All of the simulations were performed using R statistical software v.3.0.2 (R Core Team, 2013) and MLE by maximizing the likelihood of eq (1) using the optim function with the Nelder-Mead algorithm. The R code used for simulations and maximum likelihood estimation is presented in the Appendix B.

### 2.2 Assessing the Beta N-mixture Model performance

To assess the Beta N-mixture Model performance we followed three steps: (1) bias benchmark assessment, (2) comparisons with other community abundance models and (3) examples using real data. For bias benchmark assessment (section 2.2.1) we simulated 1500 data sets under the Beta N-mixture model, computed the MLEs of our model parameters each time, and then examined the distribution of the MLEs. The objective of this approach was to determine if the average of the distribution of MLEs approaches the true parameter values and if the variability around those estimates is small. In reality, data come from a much more complex process involving many variables and quantities. Therefore, in the comparison with other community abundance models (section 2.2.2), we tested the robustness of our model by simulating data from a complex, spatially explicit data-generating process, which is different from the Beta N-mixture model. For this comparison, we simulated 500 datasets and then estimated the density and detection probabilities using our model. We compared the performance of our model *vis-à-vis* a previously proposed multi-species abundance model (Yamaura *et al*., 2016). From here on, we refer to Yamaura *et al*. (2016)’s approach as the Normal N-mixture model. Finally, in the example using real data (section 2.2.3) we estimated the density of 26 species of neotropical dry forest birds using a previously unpublished dataset. The objective of this step was to illustrate the use of our model with a realistic scenario and compare the outcome of the estimates with the Normal N-mixture model.

#### 2.2.1 Bias benchmark assessment

To evaluate the bias of the Beta N-mixture model, we simulated species counts (Y_s_) in a 100-ha plot sampled using 25, point counts visited three times each. We assumed that the community was composed of 15 species, each one with a different density varying between 1 and 100 individuals/100ha (<u>λ</u>_100-_*_ha_* = 1, 2, 3, 4, 5, 7, 10, 15, 25, 40, 55, 65, 75, 85, 100). In the latter vector each value of λ_100-ha_ represents the density of a single species in the 100-ha plot. In each simulation we drew N*_sij_* individuals in each point count from a Poisson distribution with mean 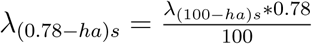.

Note that even though N*_sij_* are the realized number of individuals from the Poisson distribution with mean λ_(0_._78-*ha*)*s*_, these quantities are unobserved because the counts *y_sij_* depend on the detection process. For this simulation, as in the general specification of the model, sub-index *i* refers to the spatial replication (*i* = 1, 2, *3*, …, *r*), sub-index *j* refers to the temporal replication of the counts (*j* = 1, 2, 3, …, *t*) and the sub-index s refers to the species for which abundance is being modeled (*s* = 1, 2, *3*,…, *S*; see section 1.1 for definitions). We then simulated the detection process using a binomial distribution with parameters *N_sij_* and *p_s_*. We varied mean detection probability by assuming 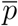 and τ = 4.5 (E[*P*] = 0.25, 0.5, 0.75; Var[P] = 0.03, 0.04, 0.03). Even though the variance seems small, the 2.5% and 97.5% quantiles of the three distributions range over a large portion of the [0,1] interval (quantiles 2.5 and 97.5: low = (0.01,0.68); mid = (0.1,0.89); high = (0.31,0.98)). For each type of community we simulated 500 data sets, and estimated λ_s_ 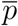 and τ using DC. To determine the number of clones required for accurate estimation of the MLEs of λ_s_ 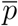 and τ we used one randomly generated data set and estimated the parameters cloning the data sequentially from 1 to 64 times (Lele *et al*., 2010). This procedure allowed us to determine an adequate number of clones to get convergence of the k^th^ posterior mean to the MLEs. We used rjags v. 4.2.0 (Plummer, 2014) with two Markov chains allowing each chain to run for 20000 generations sampling every 20 generations and discarded the first 1000 iterations. For each type of community and each simulation we estimated the relative bias 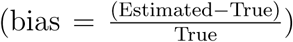 in λ_s_ 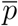 and τ.

#### 2.2.2 Comparison to other community abundance models

There are two essential differences between the Beta and Normal N-mixture models. First, Beta models treat density (λ, the mean number of individuals per sampling unit) as a fixed effect instead of random. As a result, the Normal N-mixture model has an extra hierarchy level than our model (Table 1). Both are hierarchical stochastic models where the binomial sampling model is the first hierarchy level in which the realized, but unobserved, abundances (the *N*’s) and the detection probabilities are the inner hierarchies. In both models, *N* ∼ *Poisson*(λ). The Normal N-mixture model includes an additional level and assumes that the parameters λ governing the realized abundances *N* also come from a stochastic process governed itself by hyper-parameters. In the Beta model however, *λ* does not have any hierarchy and one *λ* for each species is estimated. The second difference between our model and the Normal N-mixture model is the distributional assumption giving rise to detection probabilities. In our model *p_s_* are assumed to be 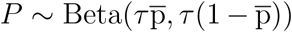 and in the Normal model, 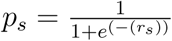 where 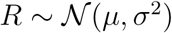, which gives a Johnson’s SB distribution between 0 and 1. Besides these two model differences, Yamaura *et al*. (2016) used a Bayesian approach to fit their hierarchical model, whereas we used the MLE method. Much discussion exists regarding the merits of each inferential approach for hierarchical models in Ecology (see for instance Lele & Dennis, 2009; Cressie *et al*., 2009). Here we limit ourselves to comparing the results from Yamaura *et al*. (2016)’s estimation approach, which is widely used as the benchmark of a known method in the literature, to our approach. Table 1 presents a comparison of the statistical models’ structures.

To compare the performance of the Normal and Beta N-mixture models we simulated 500 data sets under a spatially explicit model that had a different structure from the models evaluated (Table 1). For each data set we fitted the Normal and Beta N-mixture models and compared the posterior mean and mode estimates of the Normal N-mixture with the MLEs of the Beta N-mixture model (see Figure 2). For each simulation, we randomly drew 30 λ(_100-ha_)_s_ from a gamma distribution with parameters *α* = 0.65, β = 0.033 and excluded λ*_s_*(_100-_*_ha_*) values smaller than 1 individuals/100 ha, resulting in a community of 27 species. The gamma distribution used is the best fit of an observed species abundance distribution of a neotropical bird assemblage that was gathered using field-intensive methods (Robinson *et al*., 2000). We then randomly drew from a Poisson distribution with mean λ_(100-ha)s_, the number of individuals of the *s*^th^ species (*N_s_*) present in the 100-ha plot. We located each individual randomly across the plot and then randomly placed 25 point counts in the 100-ha plot that were separated by at least 150 meters. Finally, we obtained species-specific detection probability (*p_s_*) from a uniform distribution between 0 and 1. To obtain the counts *y_sij_*, we drew the number of individuals detected in a point count from a binomial distribution using the number of individuals in point counts *n_sij_* and the individual’s detection probability *p_s_*. We repeated the detection process three times to generate three temporal replicates of the sampling process. The R-function to simulate the described process is presented in Appendix B.

For each of the simulated data sets we estimated λ_(0.78 − *ha*)*s*_, 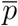 and τ under the Beta N-mixture model using ML estimation with DC (Lele *et al*., 2007). We used the variance-covariance matrix of the posterior distribution of λ(0.78 − *ha*)*s*, 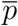 and τ To estimate Wald-type confidence intervals for each parameter (Lele *et al*., 2007, 2010). Models were built and analyzed using rjags (Plummer, 2014) with 2 chains, with iterations in each chain and retained the parameter values every 20 generations after a burn-in period of 1000 generations. After initial parameter estimation, we sampled the posterior distribution given the estimated parameters to obtain the realized values of *p_s_* given the data. For the Normal N-mixture model we performed Bayesian parameter estimation using rjags and ran the analysis using 2 chains, with iterations and retained parameters values every 20 generations after a burn-in of 10,000 generations. Even though the Normal N-mixture model is fully specified by the mean and variance of the abundance and detection processes (see Yamaura *et al*., 2016), the Beta N-mixture model has no stochastic hierarchy over λ; thus, for comparisons of the two models we retained the mean and mode of λ_(0_._78-_*_ha_*_)_*_s_*. Because *p_s_* is also a random variable with an additional level of hierarchy in the Normal N-mixture model, we also retained the mean and mode of the posterior distribution of *p_s_* resulting from Bayesian estimation. Once we obtained the estimates of λ_(0_._78-_*_ha_*_)_*_s_*, we extrapolated this estimate to λ_(100-_*_ha_*_)_*_s_* as described in sections 2.1 and 2.2.1.

#### 2.2.3 Example Using Real Data

Finally, we used a data set that consisted of 94 point counts located in three dry forest patches in Colombia. Each point count was replicated three times from January 2013 to July 2014. From this data set, we selected the understory insectivore species that forage in foliage (Karr *et al*., 1990; Parker III *et al*., 1996) to meet the requirement of the Beta N-mixture model of correlated detection probabilities among species. In total, we estimated the abundance of 26 species using both the Beta and Normal N-mixture models. We are aware that it is likely that the closed population assumption for this data set might not hold, but it is unlikely that populations of species have changed drastically from one year to another during these years. The point counts were performed in three different forest patches in the upper Magdalena Valley of Central Colombia. To maximize the sample size for abundance estimation, we aggregated the point counts into a single data set, such that the inferences of species abundances are made for the entire region instead of the particular patch. The three forest patches were separated by less than 150 km and were located within the Magdalena Valley dry forest region. Because they are in the same habitat type, the structural variables of the forest are similar and thus it is unlikely that the detection probabilities vary among patches as well as the abundance of species, allowing us to aggregate the data. Bayesian and ML estimation for the Normal and Beta N-mixture models, respectively, were performed in the same way as described previously. In order to evaluate the effect of the prior distributions on the estimates of the Normal N-mixture model, we also estimated the parameters of the Normal N-mixture using ML estimation through DC. Taking advantage of the ML estimation of the Normal and Beta N-mixture model, we further performed model selection following Ponciano *et al*. (2009)’s procedure to compute the difference in Akaike’s Information Criterion (ΔAIC) between the two models. For model selection we assumed the null model to be the Beta N-mixture model and the alternative the Normal N-mixture. 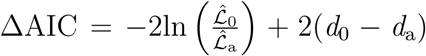, where 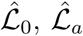 are the maximized likelihoods and *d*_0_, *d_a_* are the number of parameters of the Beta and Normal N-mixture models respectively model. Note that a ΔAIC < −2 would provide strong evidence in favor of the Beta N-mixture model, in contrast a ΔAIC > 2 would provide support in favor of the Normal N-mixture model. R code and jags models used are presented in Appendix B

## 3 Results

### 3.1 Estimation for Single Species

We found that the required minimum sample size needed for accurate estimation of the density of tropical organisms decreased when both λ and p (Figure 1) were increased. For the sample sizes evaluated, there was no combination of point counts and replicates that allowed the estimation of densities with less than 7 individuals/100ha using single-species N-mixture models (Figure A1). In the 7 ind/100ha threshold, the effort required is very high. For example, for species with a probability of detection of 0.5 the required sample size to obtain a bias lower than 0.1 is around 50 points and more than 6 replicates of each point count or around 40 point counts with more than 10 replicates (Figure 1,A1). As λ increases the sample size required to estimate appropriately the density of species decreases.

**Figure 1:**
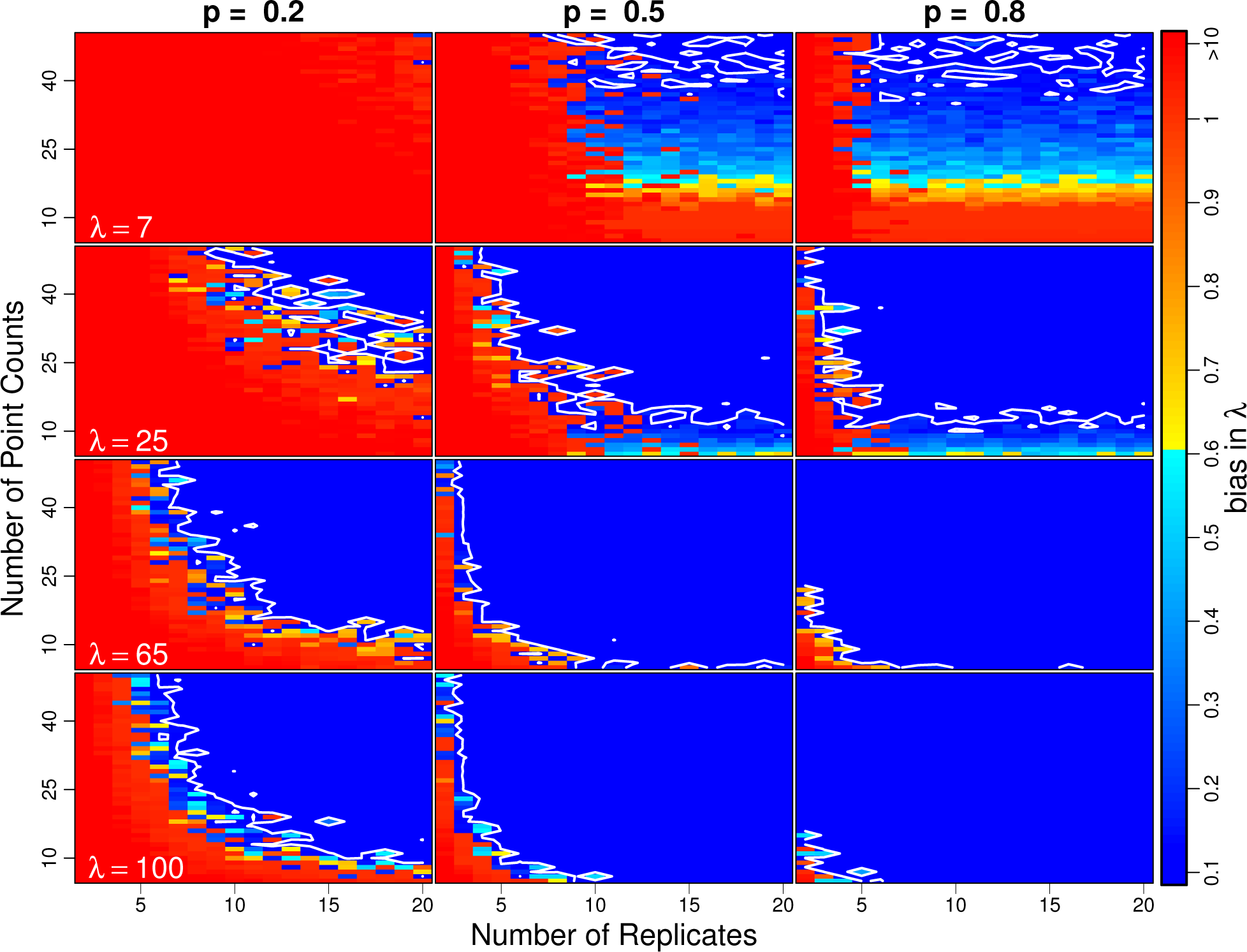
Mean bias in mean number of individuals per 100 ha λ 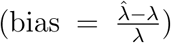 for a range of point counts, number of replicates, and true parameter values to for mid low and high abundances and detection probabilities (λ = 7, 25,65,100 and *p* = 0.2,0.5, 0.8). Colors in each panel represent the bias from low (blue) to high (red). The color scale is presented in the right. We selected a threshold for acceptable bias in estimation of abundance of 0.1 which isocline is presented as a white line in each of the panels. The results for the entire set of simulations are presented in a similar figure in appendix A

**Figure 2:**
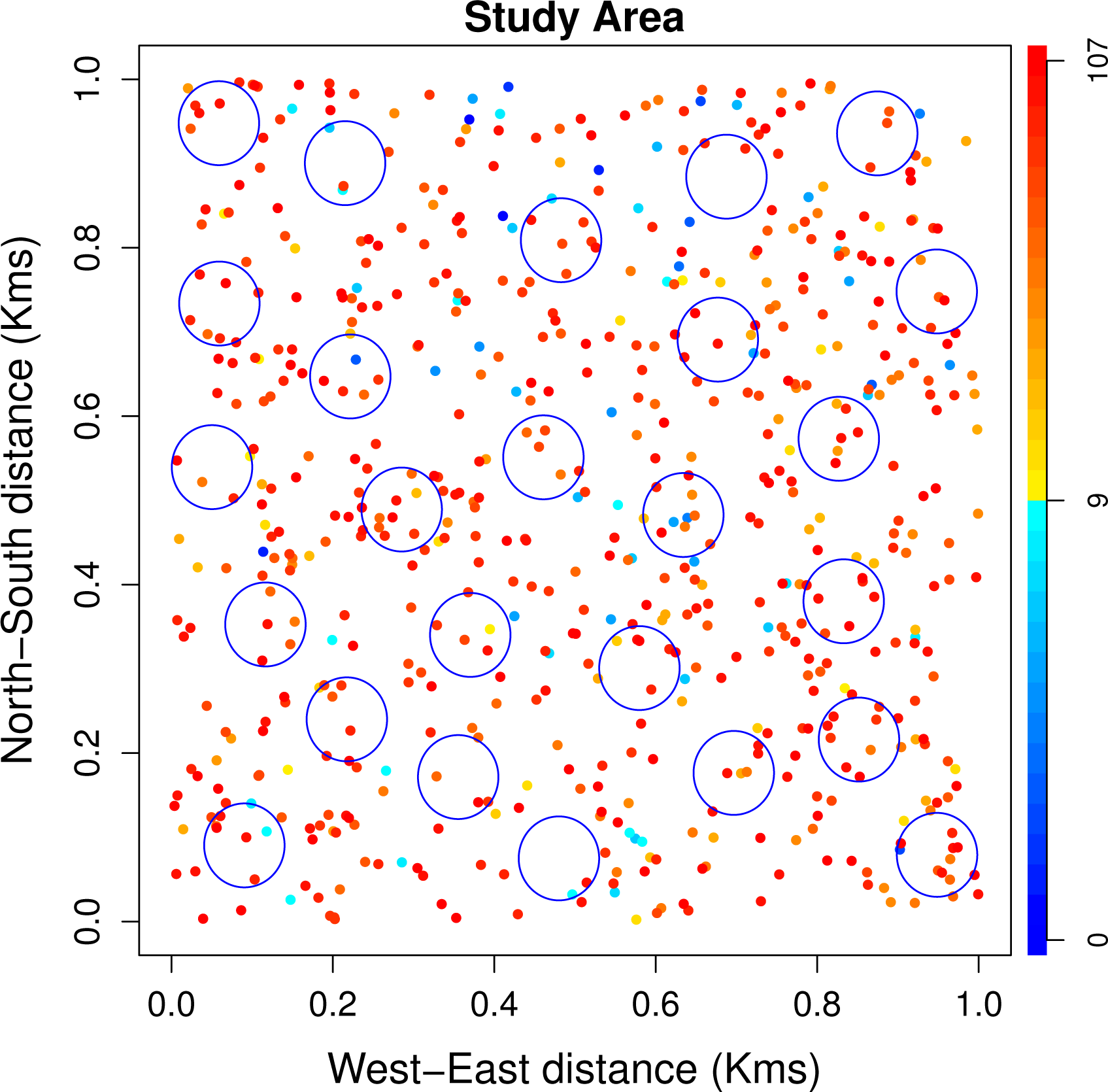
Graphic representation of the sampling design used to simulate the 500 count datasets of a community consisting of 27 species. We assumed the plot 20 be 100 ha (1 *km^2^*) and circular sampling point to be of 0.78 ha (∼ 0.008 *km*^2^). We show the true abundances in the plot represented by colors in the scale bar

### 3.2 Assessing the Beta N-mixture Model performance

#### 3.2.1 Bias Benchmark assessment

We found that the parameters of the Beta N-mixture model were fully identifiable because the relative magnitude of the first eigenvalue of the parameter variance-covariance matrix decreased very similarly at a rate of 1/*k* (*eigenvalue* = -0.07 + 1.02(1/*k*); *r*^2^ = 0.98). This result also identified that 20 clones were enough to guarantee convergence to the MLEs. The Beta model tended to slightly overestimate the density of rare species and underestimate the density of abundant species but this tendency decreased with increasing detection probability (Figure A2), as suggested by the slopes estimated by the relationship between estimated and true λ. The relationship for *p* = 0.25 was 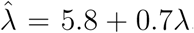, for *p* = 0.5 was 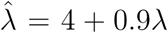 and for *p* = 0.75 was 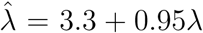,. The bias decreased (approximately) as a function of the true value of λ according to the equation bias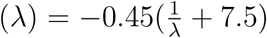 for *p* = 0.25, and bias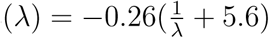 for *p* = 0.5 and bias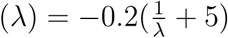 for *p* = 0.75.

Assuming that a 10% bias in the estimation is acceptable, the minimum λ that the model is able to estimate is 13 - 17 individuals/100 ha regardless of the detection probability. It is noteworthy, however, that a bias of 100% in the low-abundance end has little effects on the ecological interpretation of the estimates. Thus, if one sets bias in the abundance estimates to 100% (left hand side in the bias functions above), the model is able to predict the density of species with 3 - 5 individuals/100 ha.

The beta N-mixture model also performs well in estimating the distribution of the community’s detection probability (Figure A3). The distribution of 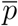 for the simulations is almost centered in the true value of p. There is a slight overestimation of 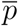 when p = 0.25 (Figure A3). The model tends to underestimate 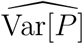, but estimates it to be similar across the different types of simulations (Figure A3).

#### 3.2.2 Comparison to other community abundance models

The beta N-mixture model performed better than the Normal model in estimating the abundance and detection probability of rare species. Whereas the posterior means and modes of the Normal model were biased towards species with abundances lower than 4 individuals/100 ha, MLEs of the Beta model were not (Figure 3). Furthermore, we found that the posterior means tended to be more biased than the posterior mode in estimating λ (Figure 3). The opposite seems to be true for the detection probabilities p. Both the posterior mode and mean underestimated *p* for rare species (Figure 4).

**Figure 3:**
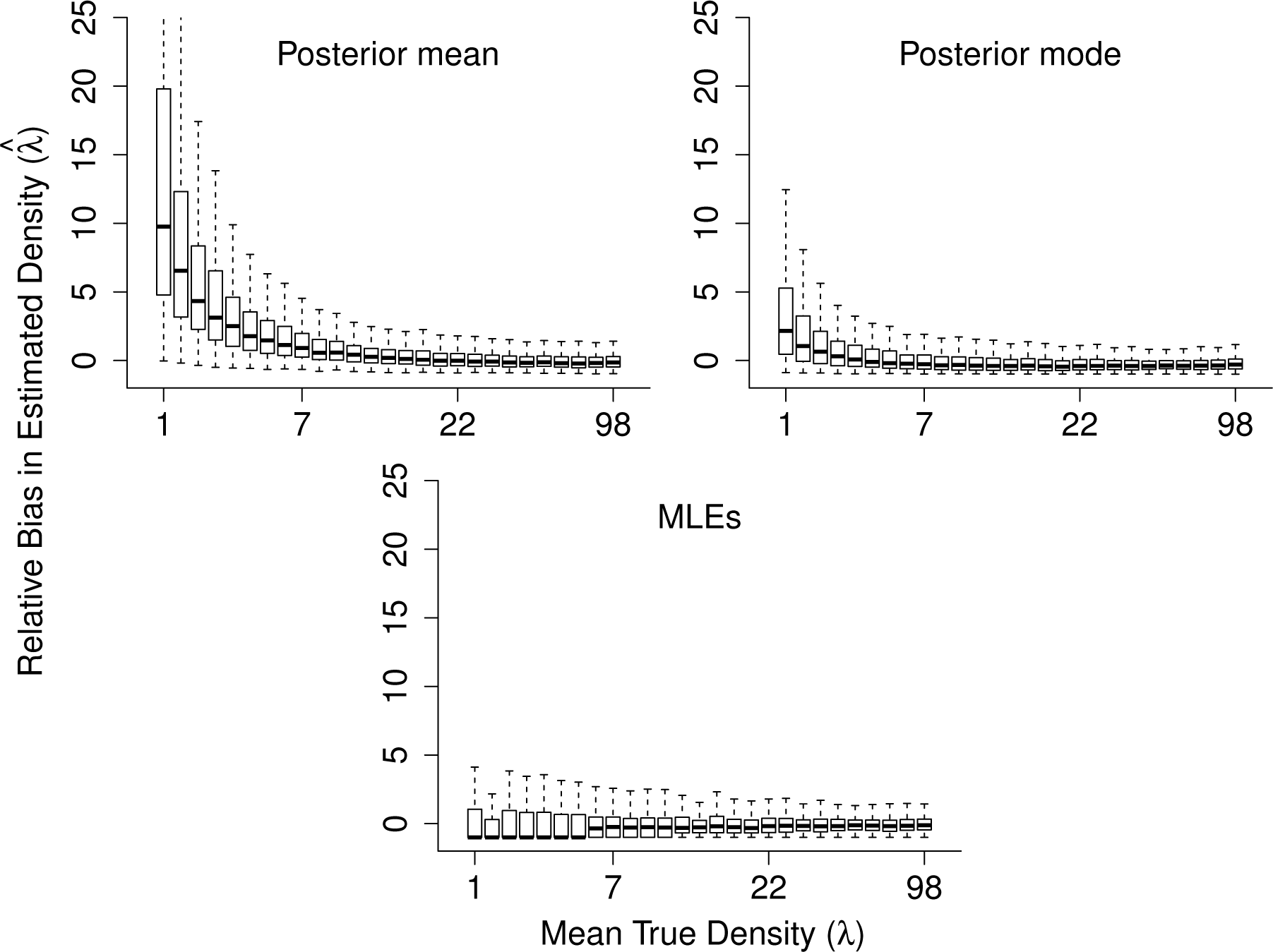
Relative bias in the estimated value of λ ((Estimate-True)/True) for both the Beta and Normal N-mixture model for 500 simulations of count data, for a community consisting of 27 species. We show the boxplots of the 500 posterior means and modes for the Normal model and the 500 Maximum Likelihood Estimates (MLEs) for the Beta model based on the same simulated data sets. The mean true abundances for each of the 27 species varied from 1 to 98 individuals/100 ha. Because there are 27 true abundances in the community the figure shows one boxplot for each species in the community.

**Figure 4:**
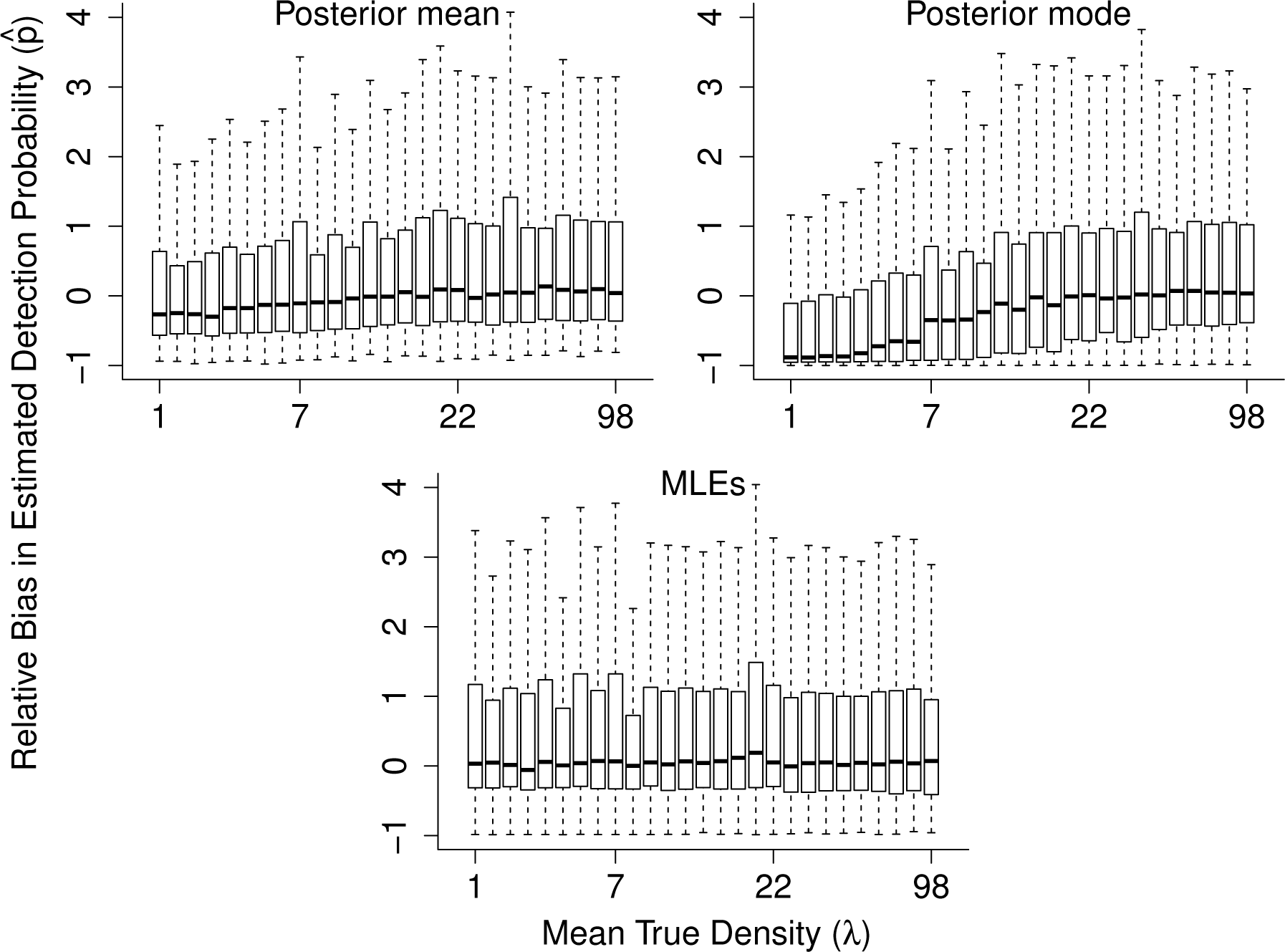
Relative bias in the estimated value of p_s_ ((Estimate-True)/True) as a function of the true abundance for both the Beta and Normal N-mixture model for 500 simulations of count data, for a community consisting of 27 species. We show the boxplots of the 500 posterior means and modes for the Normal model and the 500 Maximum Likelihood Estimates (MLEs) for the Beta model based on the same simulated data sets. The mean true abundances for each of the 27 species varies from about 1 to 98 individuals/100 ha. Because there are 27 true abundances in the community the figure shows one boxplot for each species in the community.

### 3.3 Example Using Real Data

We present the estimates of 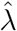 for both models in Table 2. The resulting estimates of the densities were very similar for both Beta and Normal N-mixture models (Table 2, Figure A4, Figure A5). The confidence intervals of the Beta N-mixture and Normal N-mixture overlapped for every species (Table 2). The differences in the estimates are slightly higher for rare species when estimated using the Normal N-mixture model. The Beta model estimated 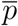 and τ = 13.5(11.9, 15). The normal model estimated *μ* = −1.22(−1.5, −1) and σ^2^ = 0.2(0.01, 0.6) or a mean detection probability of 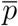 (Figure A5). The estimates of λ from the Normal N-mixture model obtained by Bayesian estimation were indistinguishable from the ones obtained from MLE (Figure A4). We found ΔAIC = −328.6 suggesting that the Beta N-mixture model is a much better fit for the counts of birds in the dry forest of the Magdalena Valley than the Normal N-mixture model.

**Table 2:** Estimates for understory insectivorous birds in the dry forest of the Magdalena Valley Colombia. Estimates are in individuals/100 ha. Det shows the number of detections of each species in the data set. We present the Upper and Lower values of the confidence interval for the Beta N-mixture model and credible interval for the Normal N-mixture model. Parker refers to the abundance category in the Parker III *et al*. (1996) database. U= Uncommon, C = Common, F= Fairly Common, F/P = Fairly common but with patchy distribution.

| Species | Det | Parker | Yamaura model |  |  | Beta model |  |  |
| --- | --- | --- | --- | --- | --- | --- | --- | --- |
|  |  |  | 97.5% | Mean | 2.5% | 97.5% | MLE | 2.5% |
| *Atalotriccus pilaris* | 83 | F | 97.3 | 145.2 | 206.1 | 71.3 | 122.8 | 174.3 |
| *Basileuterus rufifrons* | 104 | C | 146.4 | 208.6 | 300.9 | 111.2 | 204.3 | 297.3 |
| *Campylorhynchus griseus* | 7 | C | 5.0 | 14.5 | 30.1 | 0.0 | 11.2 | 22.5 |
| *Cantorchilus leucotis* | 3 | C | 2.9 | 10.3 | 24.1 | 0.0 | 8.2 | 19.5 |
| *Cnemotriccus fuscatus* | 31 | F | 39.3 | 67.0 | 110.9 | 24.3 | 67.2 | 110.2 |
| *Contopus cinereus* | 2 | F/P | 1.7 | 7.8 | 19.8 | 0.0 | 5.2 | 13.4 |
| *Cymbilaimus lineatus* | 4 | F | 4.1 | 12.9 | 28.8 | 0.0 | 11.3 | 25.0 |
| *Dromococcyx phasianellus* | 1 | U | 0.8 | 5.5 | 15.8 | 0.0 | 2.5 | 7.7 |
| *Elaenia flavogaster* | 67 | C | 107.9 | 162.8 | 260.6 | 85.7 | 192.3 | 298.8 |
| *Euscarthmus meloryphus* | 26 | C | 28.1 | 49.8 | 81.0 | 17.3 | 44.3 | 71.3 |
| *Formicivora grisea* | 172 | C | 225.4 | 315.0 | 433.1 | 172.6 | 279.0 | 385.4 |
| *Hemitriccus margaritaceiventer* | 106 | C | 104.2 | 161.6 | 231.4 | 83.6 | 124.4 | 165.1 |
| *Henicorhina leucosticta* | 28 | F | 37.7 | 65.8 | 113.6 | 20.9 | 70.9 | 121.0 |
| *Hylophilus flavipes* | 144 | C | 236.1 | 344.8 | 580.2 | 134.1 | 445.8 | 757.5 |
| *Leptopogon amaurocephalus* | 23 | F | 27.0 | 49.1 | 83.4 | 15.1 | 47.1 | 79.2 |
| *Myrmeciza longipes* | 64 | C | 81.2 | 121.6 | 178.9 | 60.1 | 111.6 | 163.1 |
| *Myrmotherula pacifica* | 1 | F | 0.8 | 5.5 | 15.4 | 0.0 | 2.5 | 7.5 |
| *Pheugopedius fasciatoventris* | 83 | F | 114.0 | 164.2 | 237.2 | 85.9 | 157.3 | 228.7 |
| *Poecilotriccus sylvia* | 69 | F | 89.2 | 135.3 | 201.7 | 61.9 | 125.4 | 189.0 |
| *Ramphocaenus melanurus* | 5 | F/P | 3.8 | 12.3 | 27.3 | 0.0 | 9.7 | 20.9 |
| *Synallaxis albescens* | 1 | C | 0.8 | 5.6 | 15.6 | 0.0 | 2.5 | 7.5 |
| *Thamnophilus atrinucha* | 93 | C | 124.1 | 177.1 | 251.6 | 91.9 | 162.7 | 233.6 |
| *Thamnophilus doliatus* | 192 | C | 269.2 | 369.7 | 516.5 | 211.2 | 345.7 | 480.2 |
| *Todirostrum cinereum* | 51 | C | 63.2 | 97.6 | 144.3 | 46.9 | 89.5 | 132.2 |
| *Tolmomyias sulphurescens* | 80 | F | 110.8 | 162.1 | 240.4 | 80.8 | 157.1 | 233.3 |
| *Troglodytes aedon* | 26 | C | 25.6 | 45.8 | 74.3 | 15.7 | 38.5 | 61.3 |

## 4 Discussion

Our results involve three major findings. First, single species N-mixture models require a high number of spatial and temporal replicates for accurate estimation of the abundance of tropical organisms (Figure 1, see also Yamaura, 2013). Second, we found that the MLEs of a wide range of abundances computed using the Beta N-mixture model have good statistical properties. Among these is a low relative bias of the parameters (*p* and λ); our approach led to unbiased estimates of the density of rare species with 1-3 individuals/100 ha (Figure 3, Figure A2). And third, we show that the MLEs of the Beta N-mixture model parameters have lower biases than the estimates provided by Yamaura *et al*. (2016)’s Normal N-mixture model (Figures 3, 4) and that in real scenarios the Beta N-mixture model fits the data better.

N-mixture models have been proven to be useful in scenarios where species are abundant (e.g. Royle, 2004; Joseph *et al*., 2009). If the objective were to estimate the abundance of a single species, our simulations provide a guide to the sampling effort required. Published databases (e.g. Parker III *et al*., 1996; Karr *et al*., 1990) include estimates of abundance of many neotropical species, which could provide general guidelines to researchers in the field about the approximate λ and the approximate sample sizes needed to correctly estimate abundance using N-mixture models.

For rare species, the solution is to use the community abundance models. Our study and Yamaura *et al*. (2016) provide two examples of how to apply the estimation of the abundance to a set of species. Our approach has the additional advantage of providing estimates with low bias even for species with low density and low detection probabilities. For example, for communities with 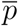 = 0.25, the mean bias for species with one individual/100 ha is around 700% (Figure A2). This number sounds extreme but it only increases the abundance from one to seven individuals/100ha having little effect on the ecological inferences drawn from the model. Furthermore, estimating the parameters of the Beta N-mixture model using a larger set of species in the community apparently corrects this bias. Our simulation under a more complex model shows that the Beta N-mixture model has almost no bias in estimating the density of species close to 1 individual/100 ha (Figure 3). The bias correction demonstrates that the larger the community, the less biased the estimates are likely to be. The latter is particularly convenient for tropical communities that are likely to have high species richness increasing the amount of information available to estimate the parameters for the entire community.

In comparison with other community abundance models (*i.e*. Yamaura *et al*., 2016), the Beta N-mixture model has lower bias in both 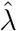 and 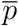. It is unknown however, why the bias toward rare species arises, because an exponential transformation of a normal distribution predicts a high number of rare species. The same scenario arises with *p* because the logit transformation of the normal distribution is more flexible than the beta distribution (Hafley & Schreuder, 1977). One explanation is that the extra level of hierarchy required by performing the transformations of the normal distribution influences estimates. Another possibility is that the prior distribution selected to perform the Bayesian estimation affects the location of the posterior means and modes. Our results, however, point to the former explanation rather than the latter, because the mean and mode of the Bayesian posterior distributions of 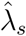 were indistinguishable from the MLEs in the real data set (Figure A4). Although in this case, prior distributions of parameters do not seem to affect the estimates, in general, prior elicitation in Bayesian analysis of hierarchical models is difficult (Lele & Dennis, 2009). In a Bayesian analysis of hierarchical models, it is important to validate the inference of these computer-intensive techniques through simulations to test the properties of posterior distributions (Dorazio, 2016; Taper & Ponciano, 2016).

One little-explored issue in the estimation of abundances using complex hierarchical models fitted *via* a Bayesian approach, is assessing when prior distributions affect the estimates of model parameters. Different uninformative priors can produce different posterior distributions that alter the inferences drawn from the model (Lele & Dennis, 2009). In particular, the use of different priors in the estimation of the probability of the detection parameter in a binomial distribution has been shown to have strong effects on the posterior distribution (Tuyl *et al*., 2008). The latter is of particular interest for community abundance estimation because the counts used to estimate abundance in community models are assumed to be binomially distributed. Strong effects from the priors might not occur in cases where the data are so extensive that the information contained in the samples overshadows the information provided by the priors. Without extensive simulations, however, it is difficult to known if this is the case. Maximum Likelihood estimation *via* DC (Lele *et al*., 2010) can be started with any prior distribution for the model parameters (as long as their support makes biological and mathematical sense) and converge to the same estimates (Lele *et al*., 2007). Also, the DC approach has the advantage that one can easily assess parameter identifiability for hierarchical models and determine when the model has too many hierarchy levels. Here, we demonstrated that all the Beta N-mixture model parameters are identifiable using Lele *et al*. (2010)’s approach.

Because our model is essentially identical to any N-mixture model, it can be adapted to any underlying distribution of abundances, although computational complications might arise in parameter estimation. Ecological inferences can be made by incorporating covariates into the abundance process as previously suggested (Joseph *et al*., 2009; Yamaura *et al*., 2011, 2012). For example, when sampling along environmental gradients, the density of species (λ) might change as a function of the gradient. In this case, λ might be estimated as a linear combination of the variables changing along the gradient. The detection process can also depend on variables influencing the overall detectability of species (Dorazio *et al*., 2013). One can assume that the detection probability distribution is a function of the functional groups or microhabitat and other species’ intrinsic characteristics that might be evolutionarily constrained (Yamaura *et al*., 2011, 2012; Ruiz-Gutiérrez *et al*., 2010). Model selection comparing models with and without abundance and detection covariates can be useful for inferring ecological mechanisms underlying the abundance of species (Joseph *et al*., 2009). In this case, ML estimation through DC is an extremely useful procedure because it allows model selection through traditional information criteria (Ponciano *et al*., 2009). In the Beta N-mixture model, the assumption of the correlated behavior can be tested by comparing it to a regular N-mixture model, and because the main difference is in the assumptions underlying detection probability, it allows us to make inferences about ecological similarity among species. Our simulations described in section 2.2.2, however, use a uniform distribution for p_s_ to generate the count data with which parameters were estimated. Such a model violates the assumption of correlated detection probabilities, but the flexibility of the beta and logit-normal distributions allow us to estimate the parameters underlying the species’ counts.

The estimates of the density of the understory insectivores of the upper Magdalena Valley show few differences between the Beta and Normal N-mixture models, except for the density of rare species (Table 2). Although the differences seem negligible at first glance, they make a big difference in the fit of the model. The ΔAIC suggested that the Beta model is by far a better fit than the Normal model for this data set, even when accounting for the larger number of parameters of the Beta model. Appropriately estimating the abundance of extremely rare species has a disproportionate effect on the fit of the models evaluated.

The abundance of more common species with higher numbers of detections in our dataset might be a little higher than in other published data sets (Karr *et al*., 1990). There are two possible reasons for this overestimation. First, when the mean detection probability of the species is low, our simulations showed that the Beta model overestimated the true abundance of species (Figure A3). Second, the data presented here comes from the dry forests of the Magdalena valley. Even though this ecosystem has lower species richness than wet forests, the biomass of the community does not change (Gomez et al. unpublished data). Populations of most species might be higher than in wet forests from which most of the abundance data for neotropical birds has been collected (Terborgh *et al*., 1990; Thiollay, 1994; Robinson *et al*., 2000; Blake, 2007).

The categorical abundance estimates from Parker III *et al*. (1996) compared to the estimates using both Beta and Normal models are similar. In particular, Table 2 shows that for most of the species that are categorized as common (C) and fairly common (F) by Parker III *et al*. (1996), the models estimate abundances to be greater than 30 individuals/100 ha. The most exciting result is the appropriate estimation of extremely rare species (*e.g*., *Dromococcyx phasianellus*), which the models accurately estimate as being rare with only 1 or 2 detections in the entire data set. These are the species that are not well estimated by the single-species models.

One of the caveats of our model is that it does not take into account unseen species (i.e., species present in the study area that are not detected during the survey). Some solutions have been suggested in a multi-species framework that would allow the estimation of at least the number of unseen species for appropriate description of the community (Dorazio & Royle, 2005; Tingley & Beissinger, 2013). Such solutions estimate the number of unseen species using occupancy modeling, but to our knowledge there are no solutions available when modeling the abundance of species. We emphasize, however, that a reasonable first step towards the objective of accurately estimating tropical species abundance distributions is to properly estimate the abundance of species that have been detected at least once.

Our simulations have pushed the limits of community abundance models by simulating species with lower yet realistic abundances than any other simulation (see Yamaura *et al*., 2016). We hope that our results encourage tropical ecologists to use community abundance hierarchical models as a means to adequately estimate the abundance of full communities. In the recent North American Ornithological congress (August 2016), two of us (JPG and SKR) participated in a discussion in which it became evident that tropical ornithologists are currently facing strong publishing challenges because abundance estimating techniques have not explicitly targeted estimation in a setting such as the tropics with very low abundances of the majority of the species and sparse counts. Unlike temperate forests, the number of species is typically very high in the tropics, but counts of individuals per species are very low. Even though our approach was developed using birds as a study system, our results suggest that it is possible to obtain reasonable estimates of the density of all of the species in a community of different taxonomic groups (e.g. mammals, insects, plants, fungi, bacteria). For example, in modeling disease ecology, it has been documented that abundance patterns in natural parasite communities is determined by host population densities, making host abundance estimation a crucial step to understand rare disease dynamics (Arneberg *et al*., 1998, e.g. ebola or avian influenza). Unbiased estimation of abundances using these hierarchical models should enable researchers to build more accurate species abundance distributions and thus seek a better understanding of the mechanisms governing biodiversity patterns (McGill *et al*., 2007).

## 5 Acknowledgements

We would like to thank the farm owners Cesar Garcia, Hacienda los Limones and Constanza Mendoza for allowing us to perform bird counts in their properties. G.Burleigh, B.Loiselle, D.Steadman, P.Shirk, associate editor and three anonymous reviewers provided useful comments for the development of the model and improvement of the manuscript. This work was supported by the National Institutes of Health Grant 1R01GM117617-01 to JKB (PI) and JMP (Co-PI).

## 6 Author Contributions

JPG and JMP conceived the ideas and designed methodology; JPG collected the data; JPG and JMP analyzed the data; JPG and JMP led the writing of the manuscript. SKR and JKB contributed critically to the drafts and gave final approval for publication.

## A Supplementary Figures

**Figure A1:**
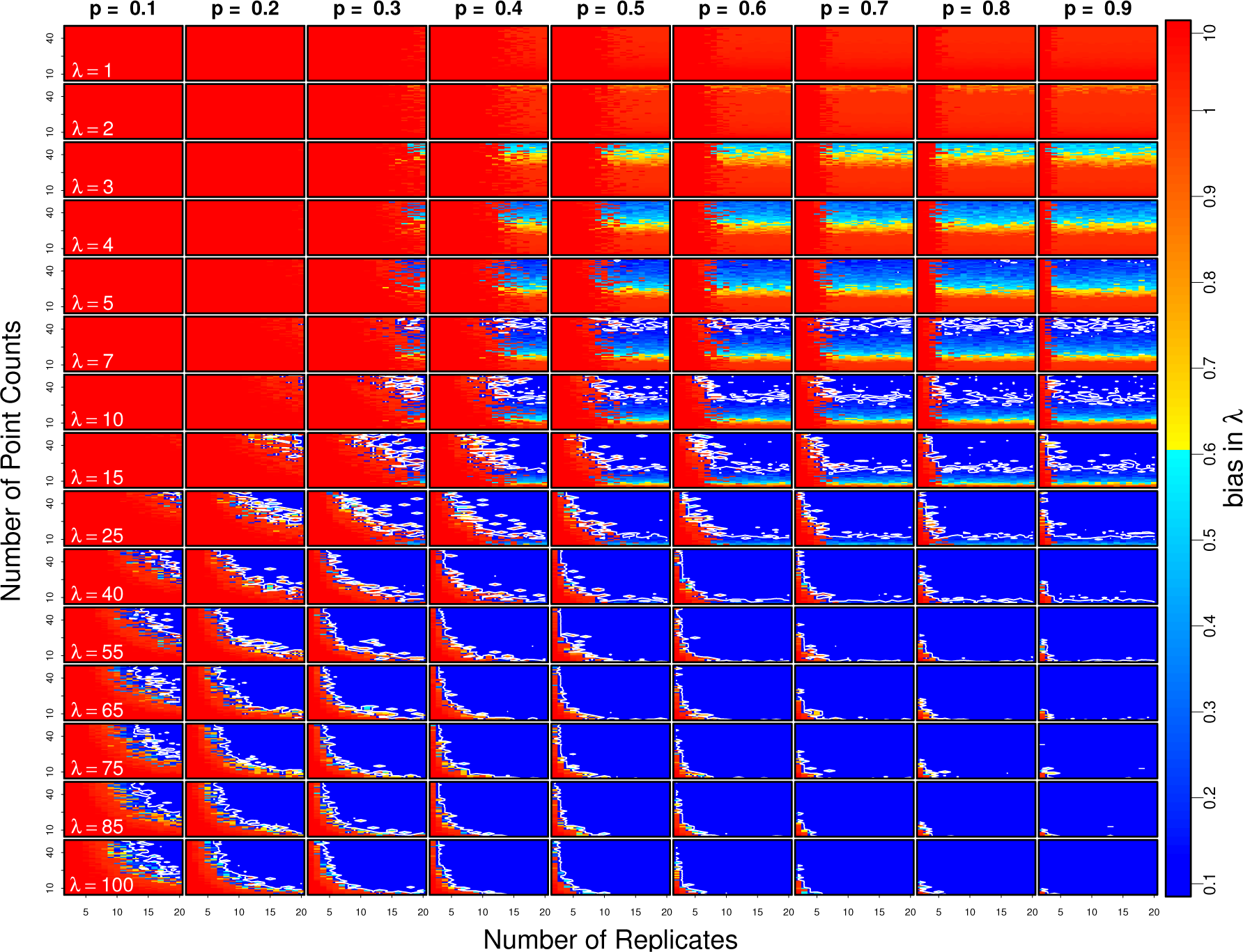
Mean bias in mean number of individuals per 100 ha λ 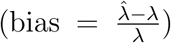 for a range of point counts, number of replicates, and true parameter values to for mid low and high abundances and detection probabilities (λ = 7, 25, 65, 100 and p = 0.2, 0.5, 0.8). Colors in each panel represent the bias from low (blue) to high (red). The color scale is presented in the right. We selected a threshold for acceptable bias in estimation of abundance of 0.1 which isocline is presented as a white line in each of the panels.

**Figure A2:**
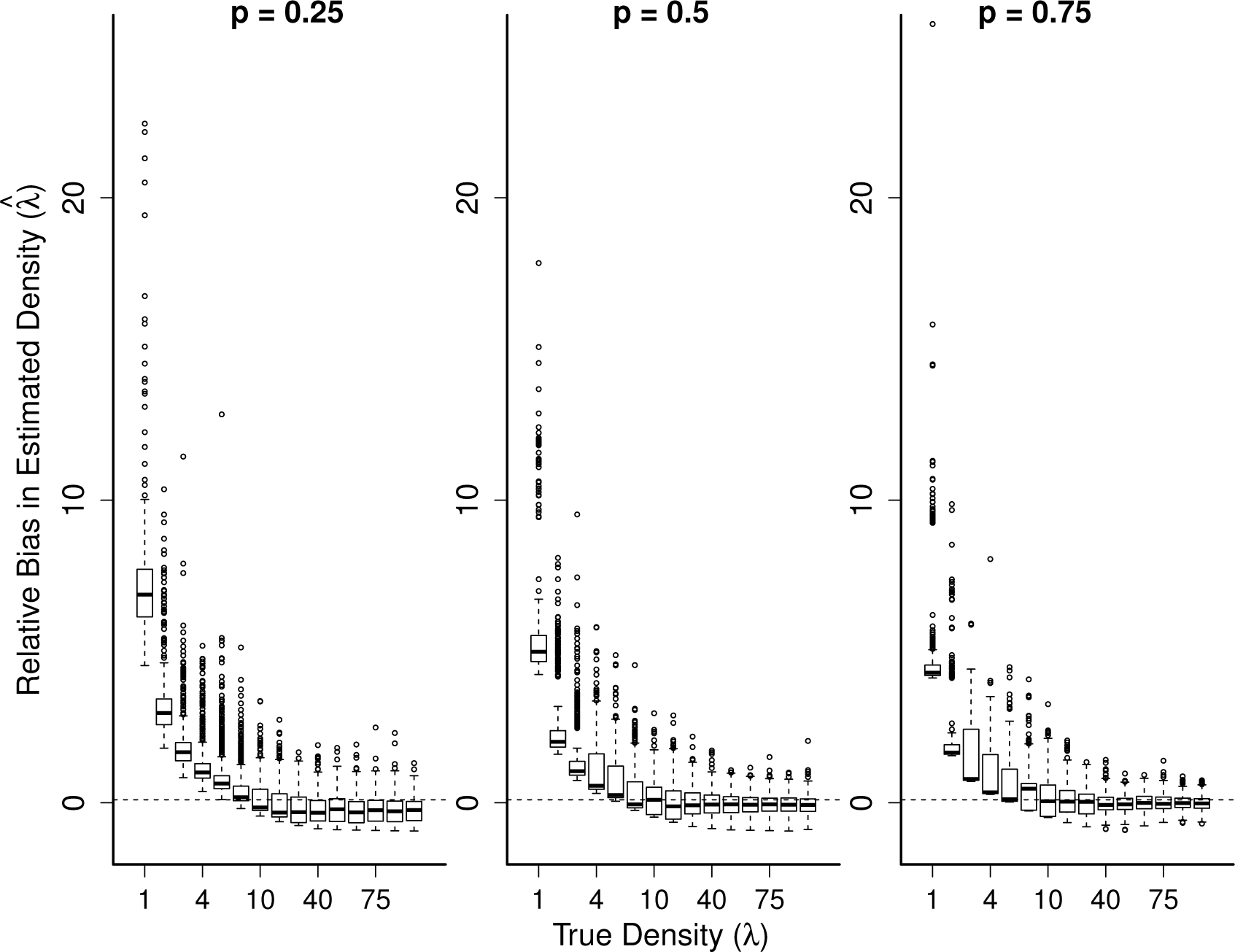
Boxplot showing the distribution of 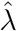 using Beta N-mixture model, showing the location of the true value of λ.

**Figure A3:**
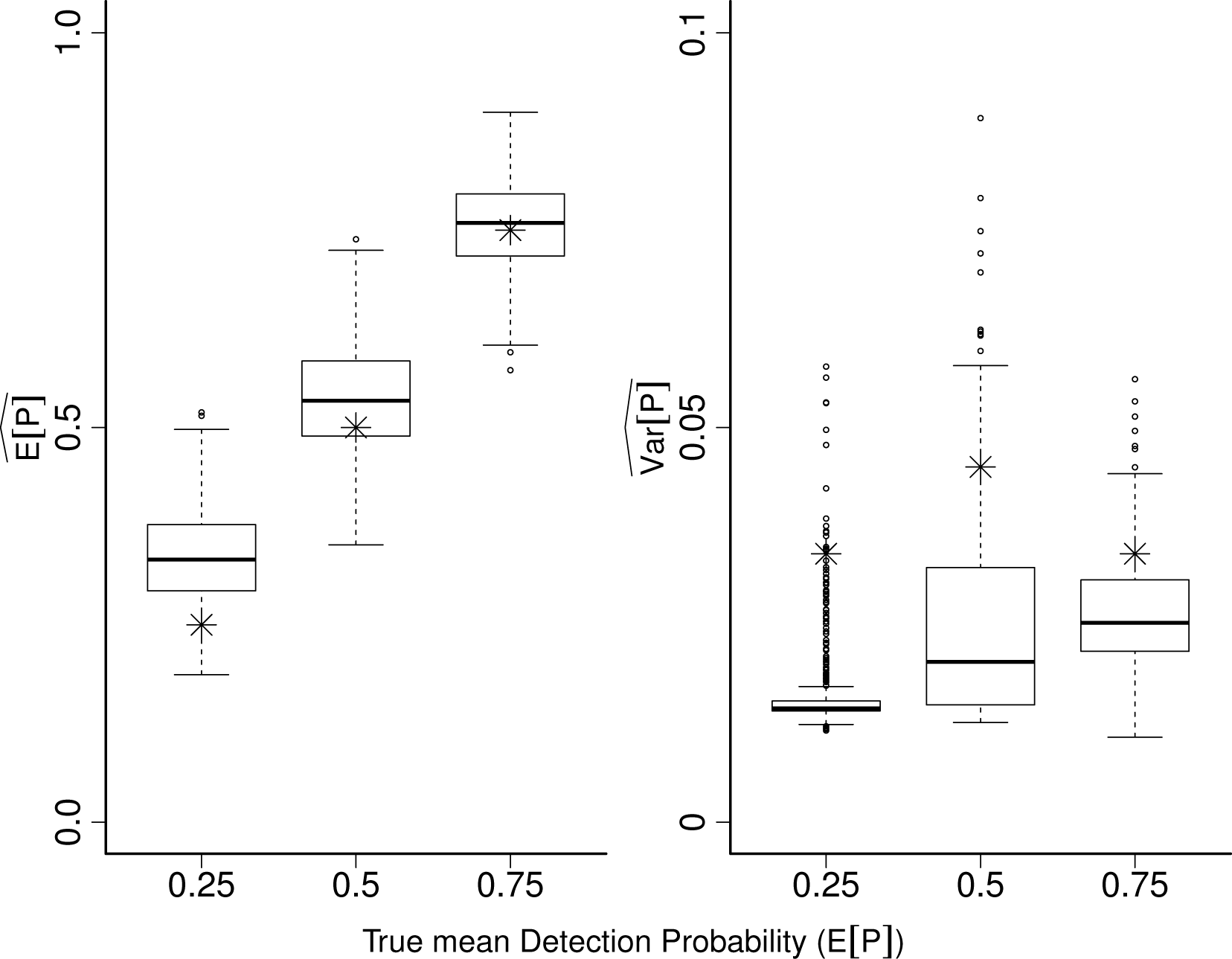
Boxplots showing the distribution of 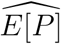 and 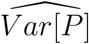 as a function of the true mean detection probability *E[P*] with which data was simulated.

**Figure A4:**
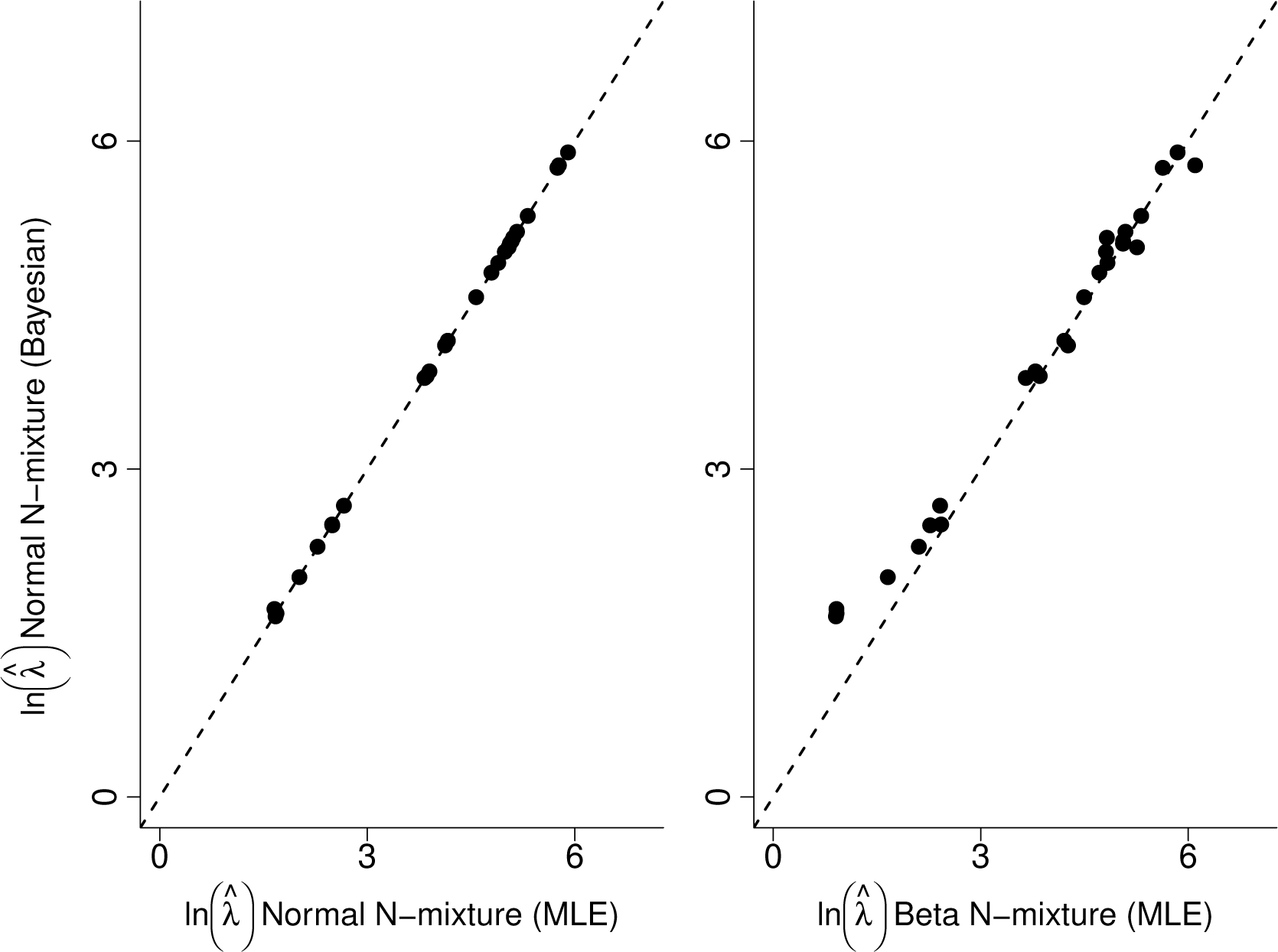
Comparison of 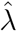 resulting from Bayesian and Maximum Likelihood estimations (MLE) of the Normal N-mixture model (left) and the estimates from the Normal and Beta N-mixture models (right)

**Figure A5:**
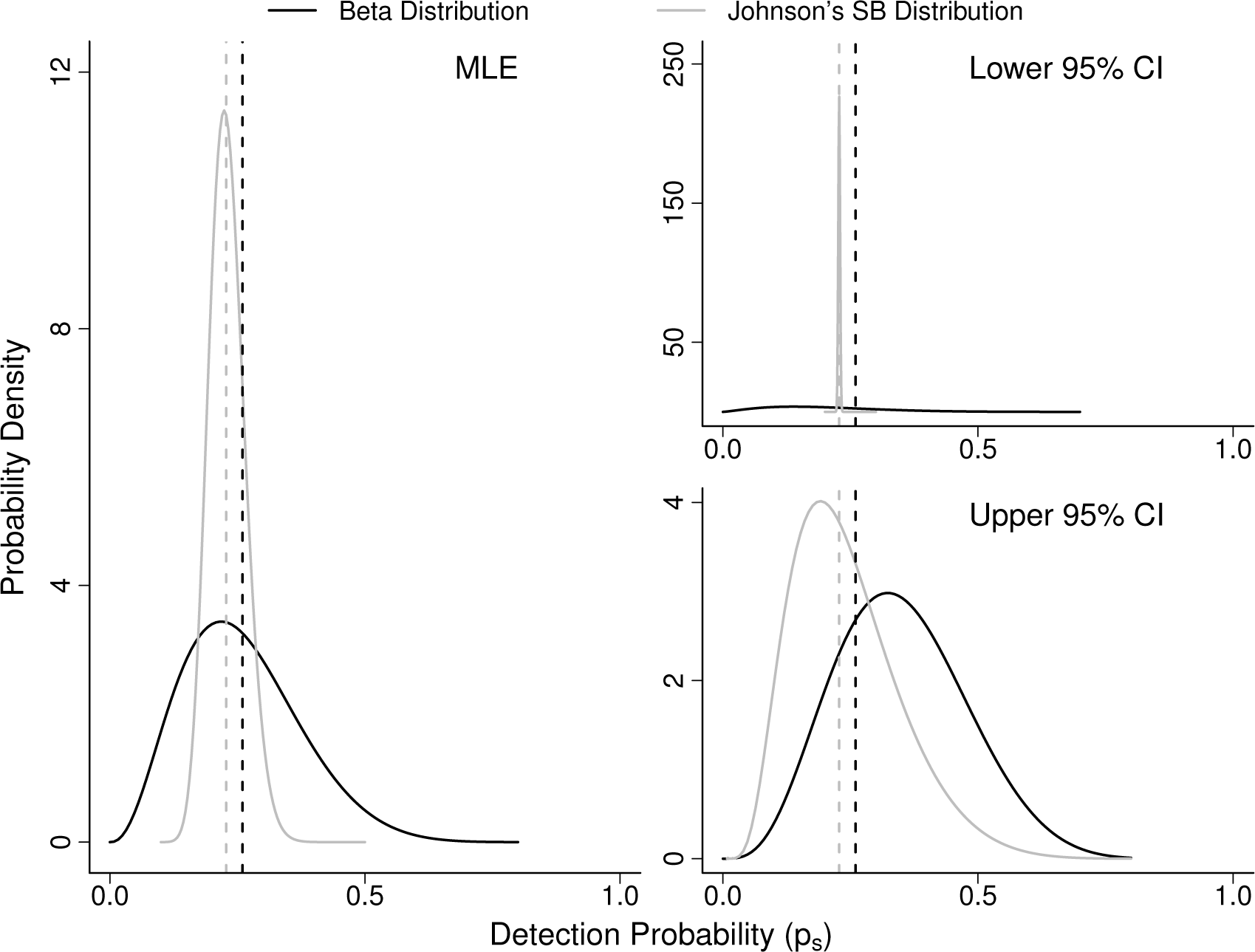
Probability distribution of the *p_s_* estimated by the Beta (black) and Normal (gray) N-mixture models for a 26 species community in the dry forest of the Magdalena River Valley in Colombia. Dotted lines represent the upper and lower curves based on the 95% confidence intervals of the parameters estimated by the models. Johnson’s SB distribution is the logit transformation of the normal distribution used to estimate detection probabilities.

## B R Code

Appendix B contains the source codes necessary for estimating abundance using the Beta and Normal N-mixture models. It is based on bugs specification of the model, R functions for abundance estimation using N-mixture model are also provided in the code. The data to the three steps of the Beta N-mixture validation are separated in different. RData files. The data sets for the 1500 simulations with hi, mid and low 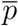 are saved in the bias.RData. The 500 data sets simulated under the complicated model used to compare the Beta and Normal N-mixture model along with the λ and p used in each simulation are saved under the comparison.RData. The real count data from the point counts performed in central Colombia are saved in the file real.RData. The entire code is saved in the Gomez_et_al_code.R from which all of the analysis of this paper can be easily replicated. The only step for which we did not save the simulated data was the bias estimation of the single species N-mixture model because of the large number of simulations performed. Using the code and function provided, however, the reader should be able to reproduce the simulations and the bias estimation.

## References

Arneberg, P., Skorping, A., Grenfell, B. & Read, A.F. (1998) Host densities as determinants of abundance in parasite communities. Proceedings of the Royal Society of London B: Biological Sciences, 265, 1283–1289.

Barnagaud, J.Y., Barbaro, L., Papaïx, J., Deconchat, M. & Brockerhoff, E.G. (2014) Habitat filtering by landscape and local forest composition in native and exotic new zealand birds. Ecology, 95, 78–87.

Bibby, C.J., Burgess, N.D., Hill, D.A. & Mustoe, S. (2000) Bird Census Techniques. Elsevier, second edition edition.

Blake, J.G. (2007) Neotropical forest bird communities: a comparison of species richness and composition at local and regional scales. The Condor, 109, 237–255.

Chandler, R.B., King, D.I., Raudales, R., Trubey, R., Chandler, C. & Arce Chávez, V.J. (2013) A small-scale land-sparing approach to conserving biological diversity in tropical agricultural landscapes. Conservation Biology, 27, 785–795.

Cressie, N., Calder, C.A., Clark, J.S., Hoef, J.M.V. & Wikle, C.K. (2009) Accounting for uncertainty in ecological analysis: the strengths and limitations of hierarchical statistical modeling. Ecological Applications, 19, 553–570.

de Valpine, P. (2012) Frequentist analysis of hierarchical models for population dynamics and demographic data. Journal of Ornithology, 152, 393–408.

Denes, F.V., Silveira, L.F. & Beissinger, S.R. (2015) Estimating abundance of un-marked animal populations: accounting for imperfect detection and other sources of zero inflation. Methods in Ecology and Evolution, 6, 543–556.

Dorazio, R.M. (2016) Bayesian data analysis in population ecology: motivations, methods, and benefits. Population Ecology, 58, 31–44.

Dorazio, R.M., Martin, J. & Edwards, H.H. (2013) Estimating abundance while accounting for rarity, correlated behavior, and other sources of variation in counts. Ecology, 94, 1472–1478.

Dorazio, R.M. & Royle, J.A. (2005) Estimating size and composition of biological communities by modeling the occurrence of species. Journal of the American Statistical Association, 100, 389–398.

Hafley, W. & Schreuder, H. (1977) Statistical distributions for fitting diameter and height data in even-aged stands. Canadian Journal of Forest Research, 7, 481–487.

Hubbell, S.P. (2001) The unified neutral theory of biodiversity and biogeography, volume 32. Princeton University Press, Princeton, NY.

Hutto, R.L., Pletschet, S.M. & Hendricks, P. (1986) A fixed-radius point count method for nonbreeding and breeding season use. The Auk, 103, 593–602.

Iknayan, K.J., Tingley, M.W., Furnas, B.J. & Beissinger, S.R. (2014) Detecting diversity: emerging methods to estimate species diversity. Trends in ecology & evolution, 29, 97–106.

Joseph, L.N., Elkin, C., Martin, T.G. & Possingham, H.P. (2009) Modeling abundance using n-mixture models: the importance of considering ecological mechanisms. Ecological Applications, 19, 631–642.

Karr, J.R., Robinson, S.K., Blake, J.G., Bierregaard Jr, R.O. & Gentry, A. (1990) Birds of four neotropical forests. A.H. Gentry, ed., Four neotropical rainforests, pp. 237–269. Yale University Press New Haven, Connecticut.

Leibold, M.A., Holyoak, M., Mouquet, N., Amarasekare, P., Chase, J.M., Hoopes, M.F., Holt, R.D., Shurin, J.B., Law, R., Tilman, D. et al. (2004) The metacommunity concept: a framework for multi-scale community ecology. Ecology letters, 7, 601–613.

Lele, S.R. & Dennis, B. (2009) Bayesian methods for hierarchical models: are ecologists making a faustian bargain. Ecological Applications, 19, 581–584.

Lele, S.R., Dennis, B. & Lutscher, F. (2007) Data cloning: easy maximum likelihood estimation for complex ecological models using bayesian markov chain monte carlo methods. Ecology letters, 10, 551–563.

Lele, S.R., Nadeem, K. & Schmuland, B. (2010) Estimability and likelihood inference for generalized linear mixed models using data cloning. Journal of the American Statistical Association, 105, 1617–1625.

MacKenzie, D.I., Nichols, J.D., Lachman, G.B., Droege, S., Andrew Royle, J. & Langtimm, C.A. (2002) Estimating site occupancy rates when detection probabilities are less than one. Ecology, 83, 2248–2255.

Margules, C.R. & Pressey, R.L. (2000) Systematic conservation planning. Nature, 405, 243–253.

Martin, J., Royle, J.A., Mackenzie, D.I., Edwards, H.H., Kery, M. & Gardner, B. (2011) Accounting for non-independent detection when estimating abundance of organisms with a bayesian approach. Methods in Ecology and Evolution, 2, 595–601.

Martin, T.G., Wintle, B.A., Rhodes, J.R., Kuhnert, P.M., Field, S.A., Low-Choy, S.J., Tyre, A.J. & Possingham, H.P. (2005) Zero tolerance ecology: improving ecological inference by modeling the source of zero observations. Ecology letters, 8, 1235–1246.

McGill, B.J., Etienne, R.S., Gray, J.S., Alonso, D., Anderson, M.J., Benecha, H.K., Dornelas, M., Enquist, B.J., Green, J.L., He, F., Hurlbert, A.H., Magurran, A.E., Marquet, P.A., Maurer, B.A., Ostling, A., Soykan, C.U., Ugland, K.I. & White, E.P. (2007) Species abundance distributions: moving beyond single prediction theories to integration within an ecological framework. Ecology letters, 10, 995–1015.

Ovaskainen, O. & Soininen, J. (2011) Making more out of sparse data: hierarchical modeling of species communities. Ecology, 92, 289–295.

Parker III, T., Stotz, D. & Fitzpatrick, J. (1996) Ecological and distributional databases for neotropical birds. D. Stotz, J. Fitzpatrick, T. Parker III & D. Moskovits, eds., Neotrpical birds: ecology and conservation. University of Chicago Press, Chicago.

Plummer, M. (2014) rjags: Bayesian graphical models using MCMC. R package version 3-13.

Ponciano, J.M., Burleigh, J.G., Braun, E.L. & Taper, M.L. (2012) Assessing parameter identifiability in phylogenetic models using data cloning. Systematic biology, 61, 955–972.

Ponciano, J.M., Taper, M.L., Dennis, B. & Lele, S.R. (2009) Hierarchical models in ecology: confidence intervals, hypothesis testing, and model selection using data cloning. Ecology, 90, 356–362.

R Core Team (2013) R: A Language and Environment for Statistical Computing. R Foundation for Statistical Computing, Vienna, Austria.

Robinson, W.D., Brawn, J.D. & Robinson, S.K. (2000) Forest bird community structure in central panama: influence of spatial scale and biogeography. Ecological Monographs, 70, 209–235.

Royle, J.A. (2004) N-mixture models for estimating population size from spatially replicated counts. Biometrics, 60, 108–115.

Royle, J.A. & Dorazio, R.M. (2008) Hierarchical modeling and inference in ecology: the analysis of data from populations, metapopulations and communities. Academic Press, San Diego, CA.

Ruiz-Gutiérrez, V., Zipkin, E.F. & Dhondt, A.A. (2010) Occupancy dynamics in a tropical bird community: unexpectedly high forest use by birds classified as non-forest species. Journal of Applied Ecology, 47, 621–630.

Sauer, J.R. & Link, W.A. (2002) Hierarchical modeling of population stability and species group attributes from survey data. Ecology, 83, 1743–1751.

Seabloom, E.W., Borer, E.T., Gross, K., Kendig, A.E., Lacroix, C., Mitchell, C.E., Mordecai, E.A. & Power, A.G. (2015) The community ecology of pathogens: coinfection, coexistence and community composition. Ecology letters, 18, 401–415.

Sibuya, M., Yoshimura, I. & Shimizu, R. (1964) Negative multinomial distribution. Annals of the Institute of Statistical Mathematics, 16, 409–426.

Stratford, J.A. & Robinson, W.D. (2005) Gulliver travels to the fragmented tropics: geographic variation in mechanisms of avian extinction. Frontiers in Ecology and the Environment, 3, 85–92.

Taper, M.L. & Ponciano, J.M. (2016) Evidential statistics as a statistical modern synthesis to support 21st century science. Population Ecology, 58, 9–29.

Terborgh, J., Robinson, S.K., Parker III, T.A., Munn, C.A. & Pierpont, N. (1990) Structure and organization of an amazonian forest bird community. Ecological Monographs, 60, 213–238.

Thiollay, J.M. (1994) Structure, density and rarity in an amazonian rainforest bird community. Journal of Tropical Ecology, 10, 449–481.

Tingley, M.W. & Beissinger, S.R. (2013) Cryptic loss of montane avian richness and high community turnover over 100 years. Ecology, 94, 598–609.

Tuyl, F., Gerlach, R. & Mengersen, K. (2008) A comparison of bayes–laplace, jeffreys, and other priors: The case of zero events. The American Statistician, 62, 40–44.

Yamaura, Y. (2013) Confronting imperfect detection: behavior of binomial mixture models under varying circumstances of visits, sampling sites, detectability, and abundance, in small-sample situations. Ornithological Science, 12, 73–78.

Yamaura, Y., Andrew Royle, J., Kuboi, K., Tada, T., Ikeno, S. & Makino, S. (2011) Modelling community dynamics based on species-level abundance models from detection/nondetection data. Journal of applied ecology, 48, 67–75.

Yamaura, Y., Kéry, M. & Royle, J.A. (2016) Study of biological communities subject to imperfect detection: bias and precision of community n-mixture abundance models in small-sample situations. Ecological Research, 31, 289–305.

Yamaura, Y., Royle, J.A., Shimada, N., Asanuma, S., Sato, T., Taki, H. & Makino, S. (2012) Biodiversity of man-made open habitats in an underused country: a class of multispecies abundance models for count data. Biodiversity and Conservation, 21, 1365–1380.

Zipkin, E.F., DeWan, A. & Andrew Royle, J. (2009) Impacts of forest fragmentation on species richness: a hierarchical approach to community modelling. Journal of Applied Ecology, 46, 815–822.

